# Parameter identifiability and model selection for sigmoid population growth models

**DOI:** 10.1101/2021.06.22.449499

**Authors:** Matthew J Simpson, Alexander P Browning, David J Warne, Oliver J Maclaren, Ruth E Baker

## Abstract

Sigmoid growth models, such as the logistic, Gompertz and Richards’ models, are widely used to study population dynamics ranging from microscopic populations of cancer cells, to continental-scale human populations. Fundamental questions about model selection and parameter estimation are critical if these models are to be used to make practical inferences. However, the question of parameter identifiability – whether a data set contains sufficient information to give unique or sufficiently precise parameter estimates – is often overlooked. We use a profile-likelihood approach to explore practical parameter identifiability using data describing the re-growth of hard coral. With this approach, we explore the relationship between parameter identifiability and model misspecification, finding that the logistic growth model does not suffer identifiability issues for the type of data we consider whereas the Gompertz and Richards’ models encounter practical non-identifiability issues. This analysis of parameter identifiability and model selection is important because different growth models are used within areas of the biological modelling literature without necessarily considering whether parameters are identifiable, or checking statistical assumptions underlying model adequacy. Standard practices that do not consider parameter identifiability can lead to unreliable or imprecise parameter estimates and potentially misleading mechanistic interpretations. While tools developed here focus on three standard sigmoid growth models only, our theoretical developments are applicable to any sigmoid growth model and any relevant data set. MATLAB implementations of all software are available on [GitHub](https://github.com/ProfMJSimpson/SigmoidGrowth).

## 1. Introduction

Classical sigmoid growth models play a critical role in many areas of ecology, and population biology [34]. Key features of these models are that they describe: (i) approximately exponential growth at small population densities where competition for resources is relatively weak; and, (ii) saturation effects at larger population densities owing to competition for resources, where the net growth rate decreases to zero as the population density approaches the carrying capacity density [34].

There are many mathematical models of biological growth with a range of resulting sigmoid growth curves [5, 52]. Within a continuum modelling framework, this general class of mathematical models can be written as

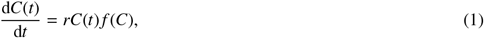

where *C*(*t*) ≥ 0 is the population density, *t* ≥ 0 is time, *r* > 0 is the intrinsic, low density growth rate, and *f*(*C*)is a crowding function that encodes information about crowding and competition effects that reduce the net growth rate as *C* increases [24]. There are many potential choices of crowding function, with typical choices of *f*(*C*) being decreasing functions, d*f*/d*C* < 0, with *f*(*K*) = 0, where *K* > 0 is some maximum carrying capacity density. Typical choices include a linear decreasing crowding function, *f*(*C*) = 1 − *C*/*K*, giving rise to the logistic growth model, or a logarithmic decreasing crowding function, *f*(*C*) = log(*K*/*C*), giving rise to the Gompertz growth model. Another choice is *f*(*C*) = 1 − (*C*/*K*)^*β*^, for some constant *β* > 0, which gives rise to the Richards’ growth model. Key features of the logistic, Gompertz and Richards’ growth models are summarised in Figures 1–2. Note that the interpretation of *r* depends upon the choice of *f*(*C*), and this coupling means that we should estimate both *r* and *f*(*C*) simultaneously from data, rather than relying on estimating *r* from low-density data and then separately estimating *f*(*C*) from high-density data [5].

**Figure 1:**
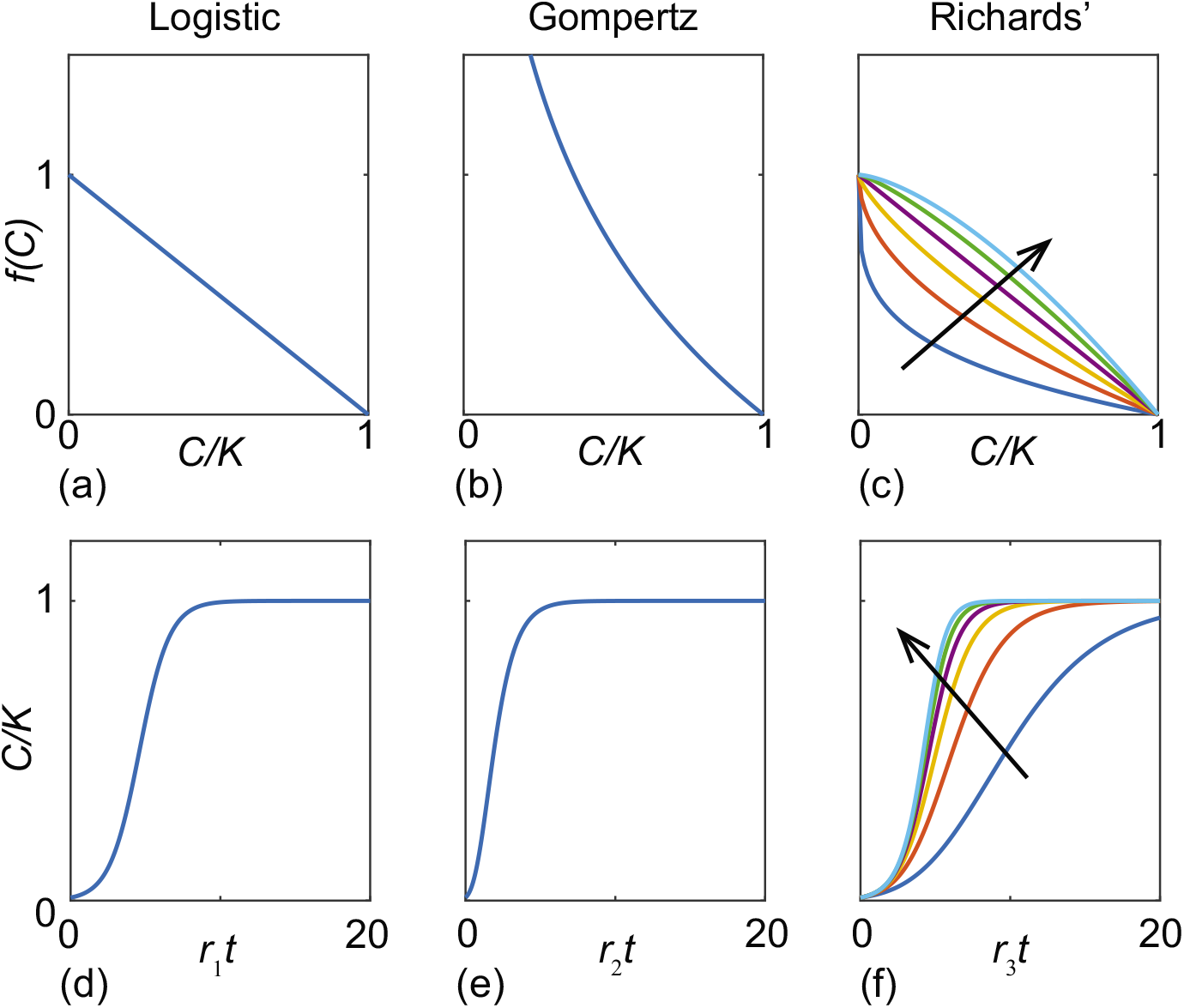
(a)-(c) *f*(*C*) as a function of *C/K* for the logistic, Gomperz and Richards’ models, as indicated. (d)-(f) Solutions of the models in (a)-(c), respectively. Curves in (c) and (f) show various profiles for *β* = 1/4, 1/2, 3/4, 1, 5/4 and 3/2, with the direction of increasing *β* indicated.

**Figure 2:**
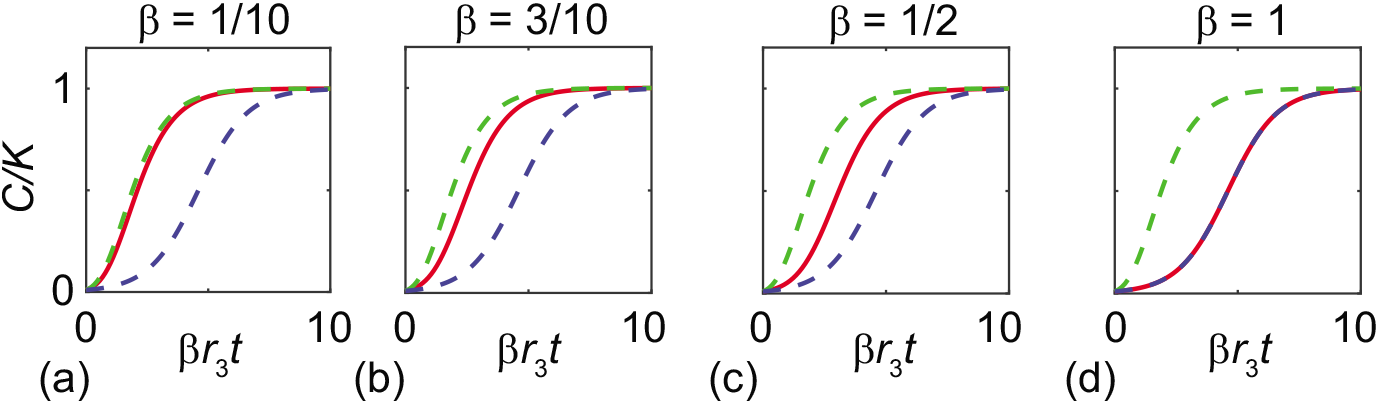
Inter-model comparison. (a)–(d) various solutions of the logistic (blue dashed), Gompertz (green dashed) and Richards’ (solid red) models with *β* = 1/10, 3/10, 1/2 and 1, as indicated, with *C*(0)/*K* = 1/100. The Gomperz solutionis shown with *r*_2_ = *βr*_3_ and the logistic solution is shown with *r*_1_ = *r*_3_.

A brief survey of the literature shows that different choices of sigmoid growth functions are routinely used automatically in different areas of application. For example, the logistic growth model is used to study a range of applications, including the dynamics of *in vitro* cell populations [8, 23, 29, 43, 55], as well as various ecological population dynamics across a massive range of scales ranging from jellyfish polyps on oyster shells [32] to continental-scale dynamics of human populations [48]. The Gompertz growth model is routinely used to study *in vivo* tumour dynamics [27, 42] as well as various ecological applications such as reef shark population dynamics [20] and dynamics of coral populations [53]. While the logistic and Gompertz models are probably the most commonly-used sigmoid growth models, various extensions have been proposed [17, 50, 52, 58].

Given the importance of sigmoid growth phenomena in ecology and biology, together with the fact that there are a number of different sigmoid growth models used to interrogate and interpret various forms of data, questions of accurate parameter estimation and model selection are crucial to ensure that appropriate mechanistic models are implemented and analysed. Indeed, the question of parameter estimation for sigmoid growth models has been addressed in the nonlinear regression literature for many years [3, 22, 39, 40], however many analyses do not address the limitations of current approaches where different discipline-specific modelling preferences are often invoked without considering other options. For example the Gompertz growth model is routinely used within the coral reef modelling community without necessarily considering other options [53, 57], while the logistic growth model is widespread within the cancer and cell biology community without necessarily considering other options [29, 43]. While such discipline-specific preference do not imply that mathematical models and their parameterisation within those disciplines are incorrect or invalid, working solely within the confines of discipline-specific choices without considering other modelling options could mean that analysts might not be using the most appropriate mathematical model to interpret data and draw accurate mechanistic conclusions.

In this work, we examine model selection for sigmoid growth models through the lens of *parameter identifiability* [2, 10, 30, 37, 38]. A model, considered as an indexed family of distributions, is formally identifiable when distinct parameter values imply distinct distributions of observations, and hence when it is possible to uniquely determine the model parameters using an infinite amount of ideal data [30, 37]. Practical identifiability involves the ability to estimate parameters to sufficient accuracy given finite, noisy data [10, 30, 37]. Methods of identifiability analysis are often used in the systems biology literature where there are many competing models available to describe similar phenomena [16, 33], and these methods provide insight into the trade-off between model complexity and data availability [7]. We also consider the often-neglected question of whether basic statistical assumptions required for the validity of identifiability analysis hold. In particular, we follow the ideas of likelihood-based frequentist inference outlined in [12, 36, 47] and partition our analysis into two types of question: (i) parameter identifiability based on the likelihood function while assuming a statistically adequate model family, and (ii) misspecification checks of basic statistical assumptions underlying model family adequacy.

There is ongoing controversy over the relative merits of various approaches to model selection in the ecological literature. For example, information-theoretic measures such as AIC [1] are increasingly used as alternatives to null hypothesis testing to select between models or carry out model averaging in ecology [9, 13, 25]. However, the underlying justifications for this trend have been criticised [19, 35, 49]. While we also consider the relationships between model complexity and model fit, we take an alternative approach, avoiding relying solely on simplistic null hypothesis testing or single number summaries such as AIC. The two components of our approach – parameter identifiability and model family adequacy – relate to fundamental statistical tasks (parameter estimation and model checking) and are easy to interpret. Furthermore, while measures such as AIC have a predictive focus and interpretation [1, 13], identifiability analysis explicitly focuses on estimating parameter values and understanding underlying mechanisms. Though prediction and estimation are related, there is often a tension between whether to ‘explain or predict’ [44].

The data we use describes concerns re-growth of hard coral cover on the Great Barrier Reef, Australia, after some external disturbance [15]. Developing our understanding of whether coral communities can re-grow after disturbances, and how long this re-growth process takes, are important questions that need to be addressed urgently as climate change-related disturbances increase in frequency [21]. If, for example, the time scale of re-growth is faster than the time between disturbances we might conclude that re-growth and recovery is likely, however if the time scale of re-growth is slower than the time between disturbances we might anticipate that re-growth and recovery is unlikely. In this work we consider a data set describing the temporal re-growth of hard coral cover on a reef near Lady Musgrave Island, Australia. Data in Figure 3 shows a typical sigmoid growth response, after some disturbance, such as a tropical cyclone, where we see apparent exponential growth for low coral cover, and the gradual reduction in net growth rate as the coral cover approaches some maximum density. Unlike small-scale laboratory experiments where it is possible to consider data from several identically prepared experiments (e.g. *in vitro* cell biology [56]), it is not possible to average this kind of reef-scale re-growth across many identically prepared experiments. Therefore, here we take the most fundamental approach and work with a single trace data set. Within the coral reef modelling community, it is typical to model this kind of sigmoid curve with a Gompertz growth model [53, 57], but in this work we objectively explore the suitability of a suite of sigmoid growth models, showing that standard approaches that ignore questions of identifiability can produce misleading results. While the focus of this work is to consider a canonical data set together with three standard choices of sigmoid growth model (logistic, Gompertz and Richards’ models), the theoretical tools and algorithms developed here can be applied to any sigmoid data set and to any continuum sigmoid growth model. One of the reasons we focus our work on the coral data is that long-term monitoring data successfully reveal the shape of the sigmoid curve, whereas applications such as cell biology often just focus on the early time data before the point of inflection [55].

**Figure 3:**
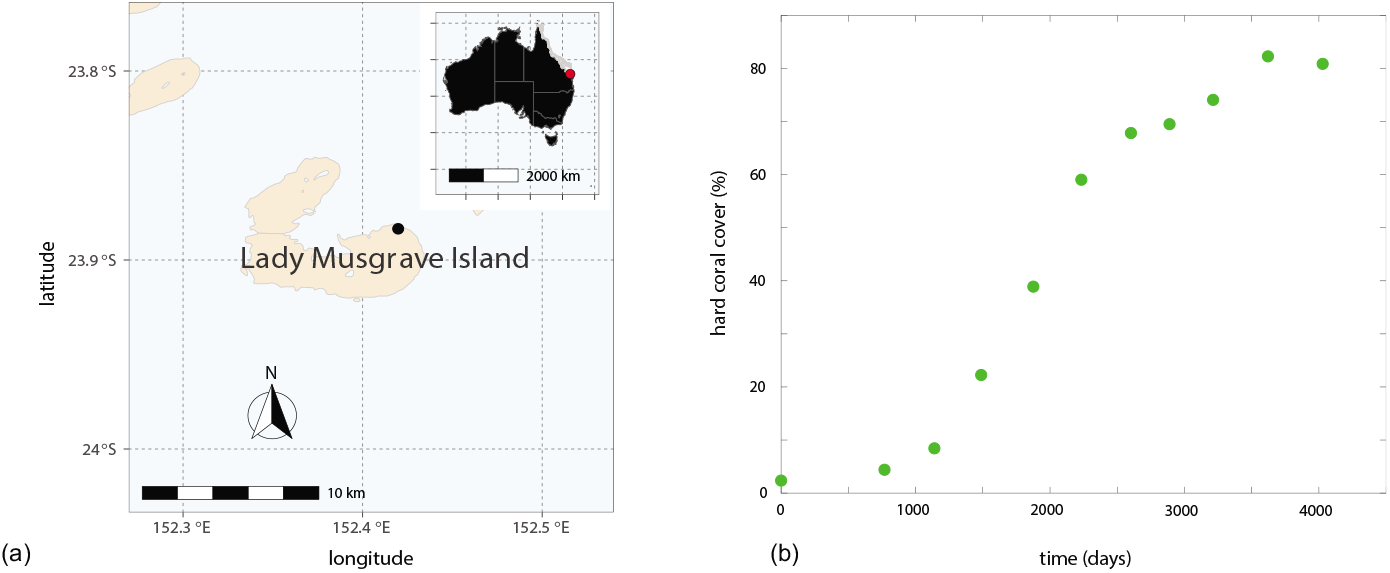
(a) Location of Lady Musgrave Island (black disc) off the East coast of Australia (inset, red disc). (b) Field data showing the % area covered by hard corals (green discs) as a function of time after some external disturbance.

## 2. Results and Discussion

### 2.1. Mathematical models

We consider three mathematical models of population growth:

–*Logistic model*

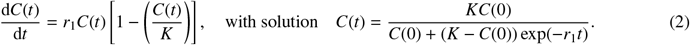 The logistic model has three free parameters, *θ* = (*r*_1_, *K, C*(0)), and a linear crowding function, *f*(*C*) = 1 – *C/K*.
–*Gompertz model*

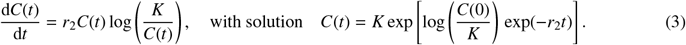 The Gompertz model has three free parameters, *θ* = (*r*_2_, *K*, *C*(0)), and a logarithmic crowding function, *f*(*C*) = log(*K/C*).
–*Richards’ model*

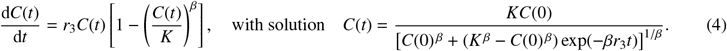 The Richards’ model has four free parameters, *θ* = (*r*_3_, *K,β, C*(0)), and a nonlinear crowding function, *f*(*C*) = 1 – (*C/K*)^*β*^, for some constant *β* > 0.

For each model the rate parameters have dimensions of 1/day, whereas *K* and *C*(0) are dimensionless, and given in terms of % of area covered by hard corals. Throughout we carefully chose our nomenclature so that certain variables (e.g. *C*(*t*), *t*) and parameters (e.g. *K*, *C*(0)) that have the same interpretation across the three models are the same for each model, whereas parameters that do not have a consistent interpretation across the models, such as the growth rates *r_m_*, for *m* = 1, 2, 3, are referred to slightly differently to make this distinction clear.

The crowding functions and solutions of each model are depicted in Figure 1 to make it clear that the question of model selection is subtle, since all three models lead to similar shaped solutions. While the three models are qualitatively similar, they are also quantitatively related as shown in Figure 2 where various solutions with different *β* are superimposed. Setting *β* = 1 in Figure 2(d) confirms that the Richards’ growth model simplifies to the logistic growth model, as expected. Further, comparing solutions in Figure 2(a)–(c) confirms that the solution of the Richards’ model approaches the solution of the Gomperz growth model, with *r*_2_ = *βr*_3_, as *β* → 0^+^. These comparisons are instructive since the limiting behaviour of the Richards’ model is trivial to establish mathematically [52], but it is not until we visually or numerically compare solutions that we gain a sense of how small *β* has to be before we can reliably approximate Richards’ growth with the simpler Gompertz growth model.

### 2.2. Parameter estimation and model checks

Data in Figure 3(b), denoted 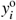, correspond to measurements at *I* discrete times, *t_i_*, for *i* = 1,2,3,…, *I*. The data, 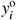, are denoted using a superscript ‘o’ to distinguish these noisy observations from their modelled counterparts, *y_i_* = *C*(*t_i_* | *θ*). To estimate *θ*, we assume the observations are noisy versions of the model solutions, 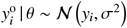, that is we assume the observation error is additive and normally distributed, with zero mean and a constant variance, *σ*^2^. The constant variance will be estimated along with the other components of *θ*. To accommodate this we include *σ* in the vector *θ* for each model.

We take a likelihood-based approach to parameter inference and uncertainty quantification given a model family. Given a time series of observations, and the above noise assumptions, the log-likelihood function is

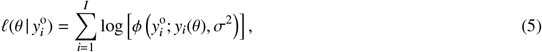

where *ϕ*(*x*; *μ*, *σ*^2^) denotes a Gaussian probability density function with mean *μ* and variance *σ*^2^. We apply maximum likelihood estimation (MLE) to obtain a best fit set of parameters, 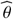. The MLE is given by

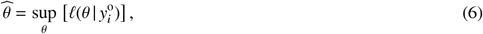

subject to bound constraints: *r_m_* > 0 for *m* = 1,2,3, *K* > 0, *C*(0) > 0, *β* > 0 and *σ* > 0. A numerical approximation of 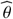 is computed using fmincon in MATLAB [31].

Given our estimates of 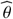, we evaluate each model at the MLE, 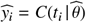 for *i* = 1,2,3,…, *I*, and calculate the scaled-residuals

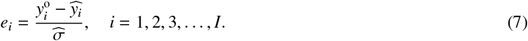

Figure 4(a)–(c) compares the data with each model evaluated at the MLE. Scaled residuals are given in Figure 4(e)–(f), and Table 1 summarises 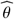 for each model. Visual checks and the Durbin-Watson test suggest that the scaled residuals are relatively uncorrelated in all cases.

**Figure 4:**
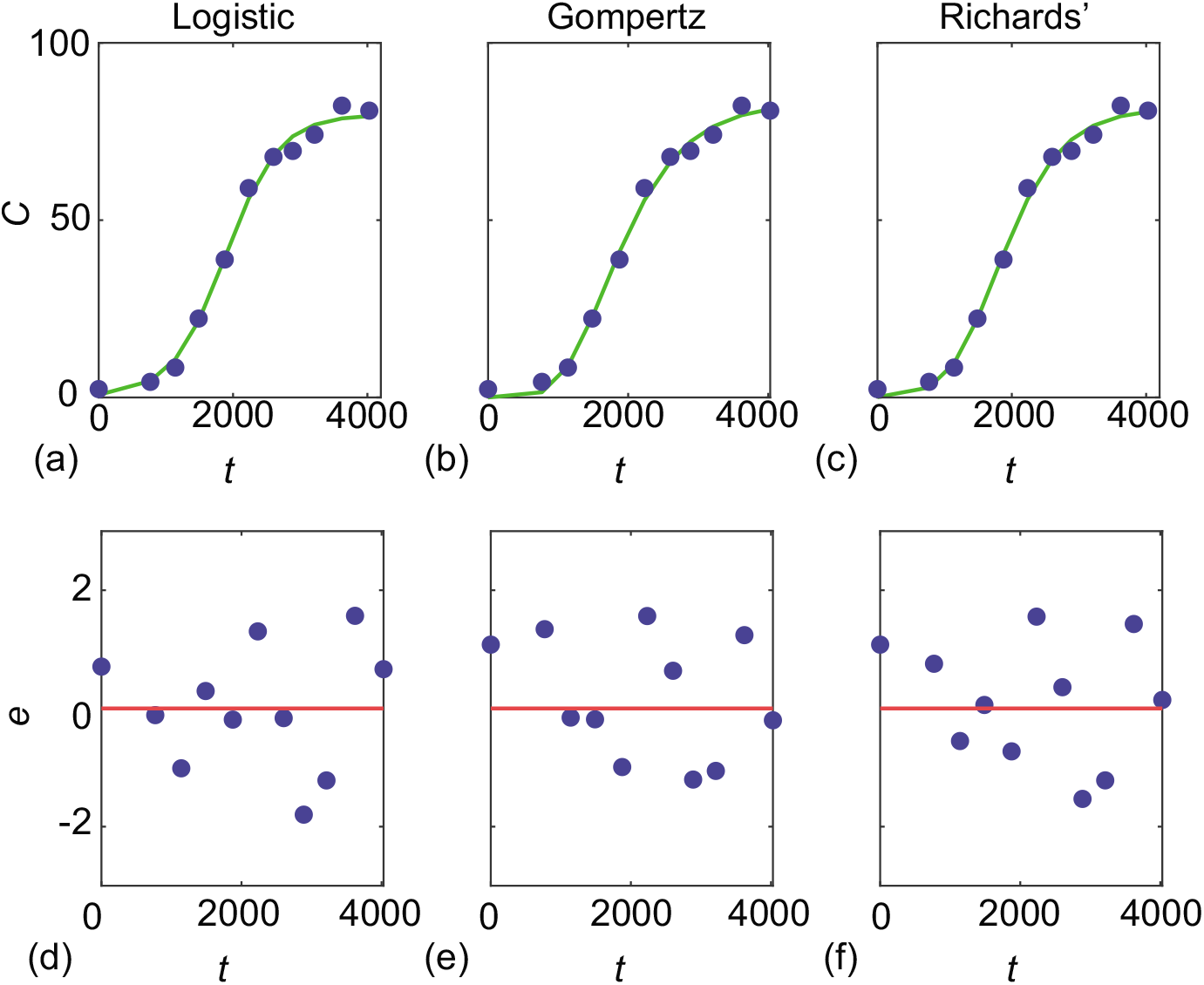
(a)-(c) Observed data (blue discs) superimposed with MLE solutions of the logistic, Gompertz and Richards’ models. (d)-(f) scaled residuals for each model. The Durbin-Watson test yields *DW* = 1.7624 (*p* = 0.3410), *DW* = 1.9894 (*p* =0.4927), and *DW* = 2.0073 (*p* = 0.5050), for the logistic, Gompertz and Richards’ models, respectively.

**Table 1:** MLE parameter estimates with 95% confidence intervals are given in parentheses.

| | Logistic ($m = 1$) | Gompertz ($m = 2$) | Richards' ($m = 3$) |
| --- | --- | --- | --- |
| $\widehat{r}_m$ | $2.5 \times 10^{-3}$ [$2.1 \times 10^{-3}, 3.0 \times 10^{-3}$] | $1.6 \times 10^{-3}$ [$1.3 \times 10^{-3}, 1.9 \times 10^{-3}$] | $5.5 \times 10^{-3}$ [$2.1 \times 10^{-3}, 1.0 \times 10^{-2}$] |
| $\widehat{K}$ | 80 [77, 83] | 83 [79, 88] | 82 [78, 87] |
| $\widehat{C(0)}$ | 0.7 [0.3, 1.4] | $1.1 \times 10^{-4}$ [$0, 1.1 \times 10^{-2}$] | $9.4 \times 10^{-2}$ [0, 1.0] |
| $\widehat{\beta}$ | - | - | 0.3 [0.1, 1.0] |
| $\widehat{\sigma}$ | 2.3 [1.6, 3.7] | 2.7 [1.5, 3.5] | 2.1 [1.4, 3.4] |

The quality of the fit to the data is excellent for each model in Figure 4(a)–(c), and visualising the scaled residuals in Figure 4(d)–(f) is consistent with the standard assumption that the scaled residuals are independent and identically distributed. The apparent uncorrelated structure of these residuals is consistent with the Durbin-Watson test. Here, simply visualising the scaled residuals confirms that standard statistical assumptions inherent in our estimation of 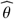 are reasonable. However, this insightful step is often overlooked, and we will return to it later.

At this point, confronted with the data-model comparisons in Figure 4 it is not obvious which model is preferable, but it is worthwhile noting some of the quantitative consequences of working with these three models. Estimates of 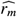 provide biological insight since the reciprocal of the growth rate, 1/*r_m_* for *m* = 1, 2, 3, provides an estimate of the time scale of re-growth to be 400, 625, and 183 days for the logistic, Gompertz and Richards’ models, respectively. While these estimates are all of the same order of magnitude, the range of 183–625 days is relatively broad if we wanted to estimate of how long we would have to wait to observe the re-growth process. If we had taken the more standard approach of interpreting this data with just one model, we would not have any insight into this variability and so this simple exercise demonstrates the importance of addressing model selection, which we now explore through the lens of parameter identifiability.

### 2.3. Identifiability analysis

In the first instance we consider the *structural identifiability* of the three models in the hypothetical situation where we have access to and infinite amount of ideal, noise-free data [16, 33]. GenSSI software [11, 28] confirms that all three sigmoid growth models are structurally identifiable. This result is very insightful since it means that identifiability issues relate to the question of *practical identifiability* which deals with the more usual scenario of working with finite, noisy data, as in Figure 3(b) [37, 38, 45].

We use a profile likelihood-based approach to explore practical identifiability. In all cases we work with a normalised log-likelihood function

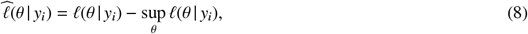

which we consider as a function of *θ* for fixed data, *y_i_*, *i* = 1,2,3,…, *I*.

We assume the full parameter *θ* can be partitioned into an *interest* parameter *ψ* and *nuisance* parameter *λ* so that we write *θ* = (*ψ, λ*). Given a set of data, *y_i_*, the profile log-likelihood for the interest parameter ψcan be written as

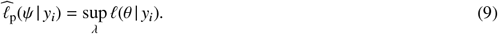

In Equation (9) *λ* is optimised out for each value of *ψ*, and this implicitly defines a function *λ*\*(*ψ*) of optimal *λ* values for each value of *ψ*. For example, in the logistic growth model with *θ* = (*r*_1_, *K*, *C*(0), *σ*) we may consider the growth rate as the interest parameter, so that *ψ*(*θ*) = *r*_1_. The remaining parameters are nuisance parameters, *λ*(*θ*) =(*K*, *C*(0), *σ*), so that

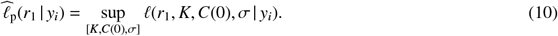

In all cases we implement this optimisation using the fmincon function in MATLAB with the same bound constraints used to calculate 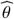 [31]. For each value of the interest parameter, taken over a sufficiently fine grid, the nuisance parameter is optimised out and the previous optimal value is used as the starting estimate for the next optimisation problem. Uniformly spaced grids of 100 points, defined on problem-specific intervals, are reported throughout.

Assuming an adequate model family, the likelihood function represents the information about *θ* contained in 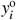, and the relative likelihood for different values of *θ* indicates the relative evidence for these parameter values [47]. A flat profile indicates non-identifiability and the degree of curvature is related to the inferential precision [36, 47]. We form approximate likelihood-based confidence intervals by choosing a threshold-relative profile log-likelihood value; for univariate profiles thresholds of −1.92 correspond to approximate 95% confidence intervals [36, 41], and the points of intersection are determined using linear interpolation. Following the likelihood-based frequentist ideas of [12, 47], we supplement the likelihood inferences about parameters given a model structure by checks of model family adequacy, here based on residual checks. We use both visual inspection and the classical Durbin-Watson test [14] to assess residual correlation, noting that we have a simple trend model and relatively small sample sizes.

Figure 5 shows various univariate profiles for each mathematical model. Each profile is superimposed with a vertical line at the MLE, and a horizontal line at −1.92 so we can visualise and calculate the width of the confidence intervals reported in Table 1. Univariate profiles in Figure 5 provide information about the practical identifiability of each model with the same data. Results for the logistic growth model indicate that each parameter is identifiable with relatively narrow profiles (Table 1). In contrast, the Gompertz growth model results show that *r*_2_ and *K* are identifiable, but *C*(0) shows signs of practical non-identifiability. In particular, the MLE and upper confidence bound for *C*(0) are very near the lower edge of feasibility, and the lower confidence bound is only determined by the *C*(0) > 0 constraint (indicating that this may be at best one-sided identifiable). Profiles for the Richards’ growth model indicate that while *K* is identifiable, the remaining parameters are not since *r*_3_ is one-sided identifiable, and profiles for *C*(0) and *β* are relatively flat, with confidence intervals determined by the *a priori* bounds rather than data.

**Figure 5:**
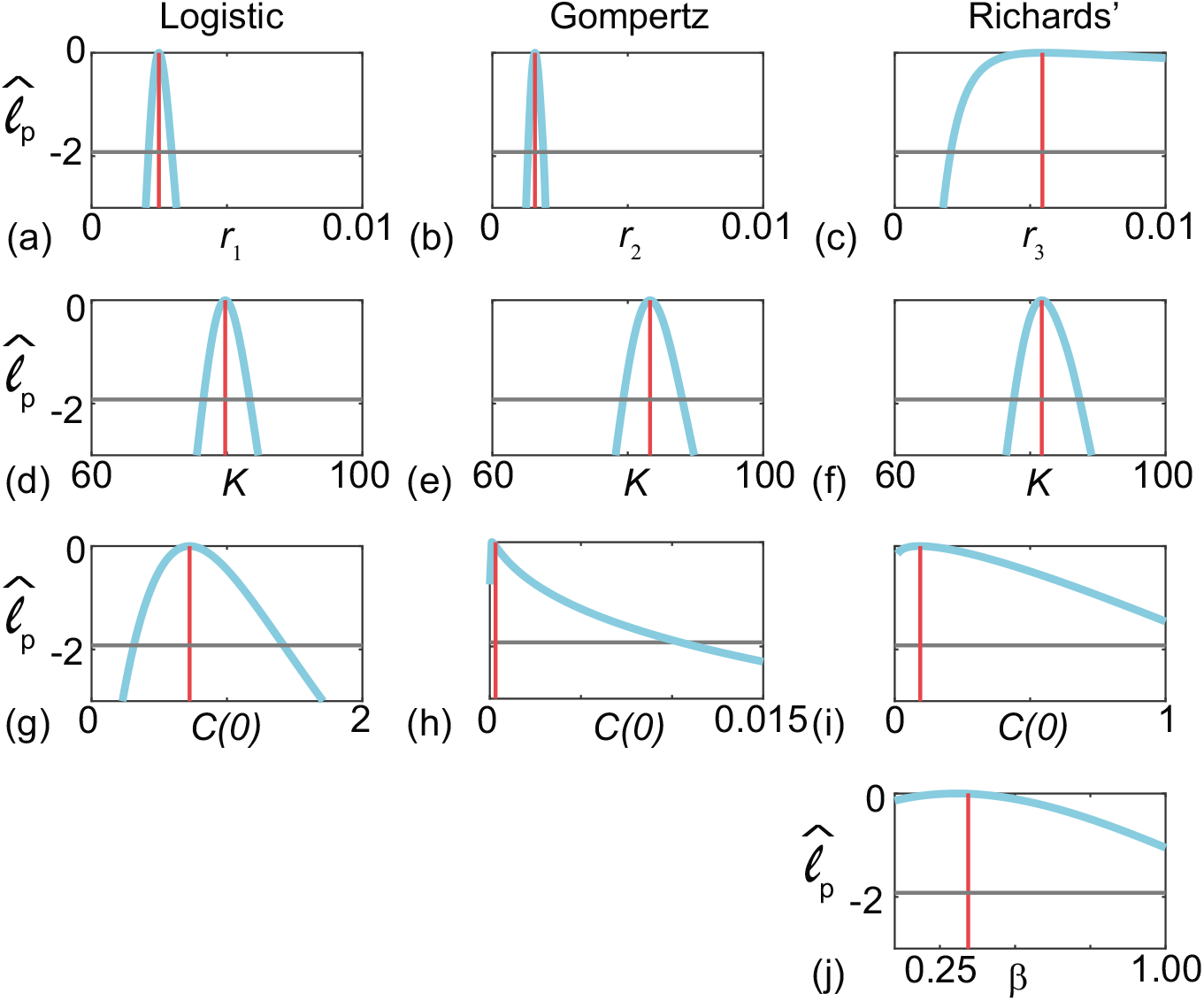
Univariate profile likelihoods for parameters in the logistic, Gompertz and Richards’ model, as indicated. In each case the profile is shown (solid blue) together with the MLE (vertical red) and a horizontal line at −1.92.

Bivariate profiles can be used to provide further insight into the identifiability issues with the Richards’ model.

In particular, we consider bivariate profile likelihoods by taking the full parameter vector, *θ* =(*r*_3_, *K*, *β*, *C*(0), *σ*), and treating *ψ*(*θ*) = (*r*_3_, *β*) and *λ*(*θ*) = (*K*, *C*(0), *σ*), allowing us to evaluate

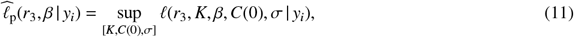

which again we compute using fmincon [31] on a uniform mesh of pairs of the interest parameter, and optimise out the nuisance parameters at each mesh point. Using uniformly-spaced grids of 100 × 100 across the interval of interest, and a threshold of −3.00 to give the approximate 95% confidence regions [36, 41] we generate the bivariate profile in Figure 6(a) where the solid grey lines define the boundary of the 95% confidence region, which, suggests there are many combinations of *β* and *r*_3_ that match the data. Finally, we now consider the product *βr*_3_ as an interest parameter, and univariate profiles for this product in Figure 6(b) suggests that our data is sufficient to obtain precise estimates of the product *βr*_3_, but does not identify *β* and *r*_3_ individually.

**Figure 6:**
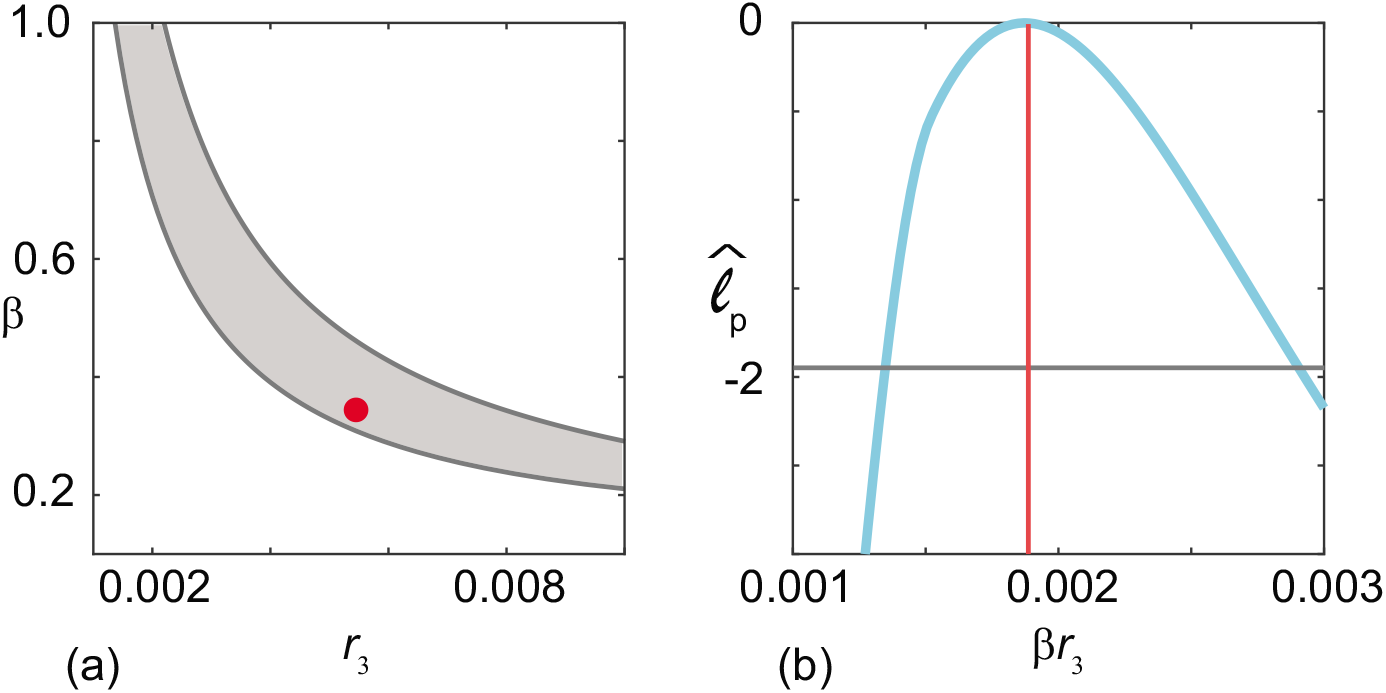
(a) Bivariate profiles for *β* and *λ*_3_. (b) Univariate profile for the product *βr*_3_. The MLE is 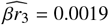 [0.0014,0.0029].

In summary, comparing the profiles between different models in Figure 5 suggests that this data leads to identifiable parameter estimates for the logistic growth model, but that the two most straightforward extensions of that model, namely the Gompertz and Richards’ models, suffer from practical identifiability issues. Therefore, we interpret these results to mean that the most reliable way to interpret this data is with the logistic growth model. While this approach to model selection using parameter identifiability is not routinely considered, it provides a straightforward, computationally-efficient means of informing quantitative distinctions between model suitability. We note, however, that difficulties with identifiability do not mean that these models are wrong, simply that they are difficult to reliably estimate parameters for, given the type of data available. This motivates our next investigation.

### 2.4. *Parameter estimation and model checks with fixed C*(0)

Our observation that *C*(0) for the Gompertz model (Figure 5(h)) is not well-identified by the data prompts us to consider a further set of results where we adopt the standard practice of estimating θunder the assumption that variability in *C*(0) is neglected, and is treated as a known quantity given by the first observation, 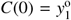 [53, 57]. Figure 7(a)–(c) compares the data with the model evaluated at the MLE with 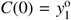, and the scaled residuals are shown in Figure7(e)–(f). A summary of the MLE estimates for each model are given in Table 2. The Durbin-Watson test indicates that the scaled residuals are more correlated when *C*(0) is fixed.

**Figure 7:**
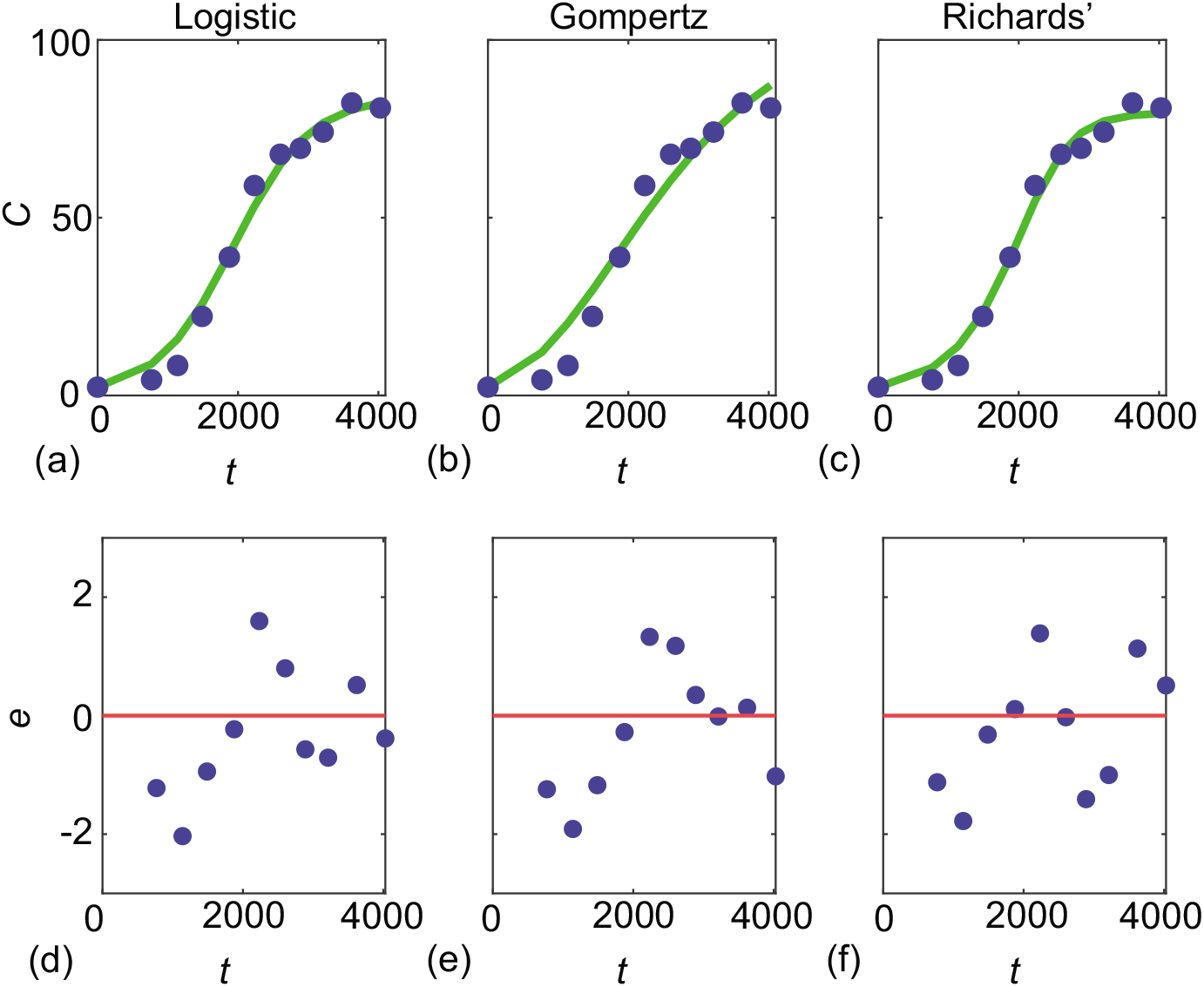
(a)-(c) Observed data (blue discs) superimposed with MLE solutions of the logistic, Gompertz and Richards’ models with 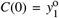. (d)-(f) scaled residuals for each model. The Durbin-Watson test yields *DW* = 1.0544 (*p* = 0.04841), *DW* = 0.6397 (*p* = 0.0045), and *DW* = 1.2919 (*p* = 0.1138), for the logistic, Gompertz and Richards’ models, respectively.

**Table 2:** Parameter estimates with fixed 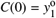, 95% confidence intervals are given in parentheses.

| | Logistic ($m = 1$) | Gompertz ($m = 2$) | Richards' ($m = 3$) |
| --- | --- | --- | --- |
| $\widehat{r}_m$ | $1.8 \times 10^{-3}$ [$1.7 \times 10^{-3}, 2.0 \times 10^{-3}$] | $7.0 \times 10^{-4}$ [$5.9 \times 10^{-4}, 9.0 \times 10^{-4}$] | $1.6 \times 10^{-3}$ [$1.5 \times 10^{-3}, 1.9 \times 10^{-3}$] |
| $\widehat{K}$ | 84 [79, 90] | 106 [88, 138] | 80 [75, 86] |
| $\widehat{\beta}$ | - | - | 1.8 [1.0, 3.9] |
| $\widehat{\sigma}$ | 3.7 [2.5, 5.9] | 6.3 [4.3, 10.2] | 3.1 [2.1, 5.0] |

The quality of the model-data match in Figure 7(a)–(c) is clearly poorer than when we simultaneously estimate *C*(0) (Figure 4), and in particular we see that the MLE solutions systematically overestimate the data at early times, and then underestimate the data at later times. These trends are clear in the scaled residuals in Figure 7(d)–(f) that are visually correlated. The Durbin-Watson test indicates that the scaled residuals are more correlated when *C*(0) is fixed. This is particularly true for the Gompertz model where we see the potential for drawing erroneous conclusions about the underlying biological mechanisms since 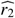 differs by an order of magnitude depending on whether we treat *C*(0) as a fixed or variable quantity. The estimate of the time scale of re-growth is 625 days in the case where we estimate *C*(0) from the data whereas it is 1429 days variability in *C*(0) is neglected. This exercise clearly illustrates the danger in adopting discipline-specific procedures without carefully considering alternative approaches.

### 2.5. Profiling misspecified models

To conclude we present some further synthetic data exploration to mimic the commonly-encountered scenario where we have sparse, noisy data generated by a relatively complicated process, which we then interpret with a simpler mathematical model. Data in Figure 8(a)–(d) show solutions of the Richards’ growth model over a range of *β* ∈ [1/10,3/2], where we consider the time interval *t* ∈ [0, *T*] where *T* is defined implicitly by *C*(*T*) = 0.999*K*. Noisy data are generated by taking 20 equally-spaced temporal observations within this interval and corrupting the data by adding normally distributed noise with zero mean and constant variance, *σ*^2^. Data in Figure 8(a)–(d) are generated by setting *r*_3_ = 0.0055, *K* = 80, *C*(0) = 1 and *σ* = 2, which are consistent with the MLE in Table 1.

**Figure 8:**
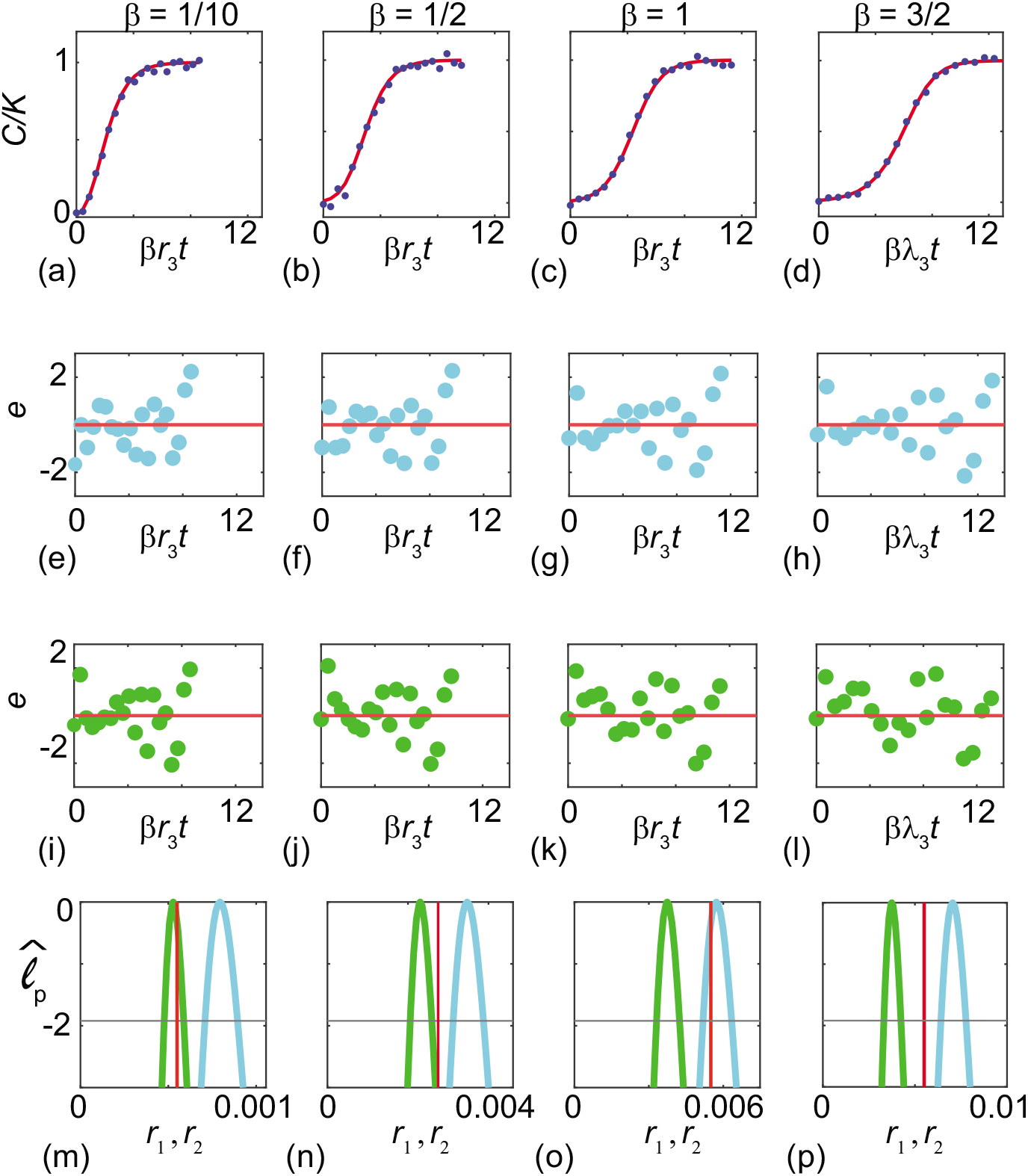
(a)–(d) Solution of the Richards’ growth curves (solid red) for the values of *β* indicated with *C*(0) = 1, *r*_3_ = 0.0055 and *K* = 80. Noisy data (blue dots) are obtained by sampling 20 equally-spaced observations in the interval 0 < *t* < *T* and adding normally distributed noise with zero mean and *σ* = 2. Here *T* is defined implicitly by *C*(*T*) = 0.999K. For each data we estimate 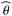 using the methods in the main paper (Figure 4, Table 1). Scaled residuals for the MLE using the logistic growth model are given in (e)–(h) and for the Gompertz growth model are given in (i)–(l). Univariate profiles for *r*_1_ (blue) and *r*_2_ (green) are given in (m)–(p). Subfigures (m)–(p) contain a red vertical line at *βr*_3_ = 0.0055*β*.

To explore how well we can interpret noisy data from the relatively complicated Richards’ growth model with the simpler logistic and Gompertz growth models we estimate 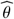 and calculate univariate profiles for *r*_1_ and *r*_2_. Scaled residuals for the logistic and Gompertz models in Figure 8(e)–(h) and Figure 8(i)–(l), respectively, where we see no obvious visual evidence of correlation. Taking the noisy data from the Richards’ growth model and constructing univariate profiles for *r*_1_ and *r*_2_ leads to well-shaped, relatively narrow profiles in Figure 8(m)–(p). These profiles again highlight certain relationships between the Richards’, logistic and Gompertz growth models established in Figure 4. In particular, for sufficiently small *β* in Figure 8(m) we see the expected relationship between the Richards’ growth model and the Gompertz growth model since the MLE for *r*_2_ is very close to the expected limiting behaviour, *r*_2_ = *βr*_3_ as *β* → 0^+^. Furthermore, not only does the profile match the expected point estimate, but we see that the width of the profile is relatively narrow, implying that it is possible to recover relatively precise and interpretable parameter estimates working with a misspecified model. Similarly, when working with the special case *β* = 1 in Figure 8(o) we see the profile for *r*_1_ for the logistic growth model is peaked close to the expected result, *r*_1_ = *r*_3_. Again, this profile relatively narrow, implying that it is possible to obtain meaningful and interpretable parameter estimates when working with a misspecified model. The remaining profiles in Figure 8(m)–(p) lead to well-shaped profiles with relatively narrow confidence intervals. Generating data from either the logistic or Gompertz models at the MLE for *r*_1_ and *r*_2_, respectively, leads to relatively uncorrelated residuals and an excellent match to the data in Figure 8(a)–(d) (not shown).

We now further explore the extent to which we are able to estimate parameters under model misspecification by repeating the calculation of 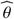 and univariate profiles for *r*_1_ and *r*_2_ using data from the Richards’ growth model by considering 30 equally-spaced values of *β* ∈ [1/10,3/2]. We repeat this exercise using both relatively noisy data with *σ* = 2, and with noise-free data, *σ* = 0. For each set of synthetic data set we generate 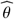 and compute the quantities 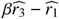 and 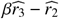, that are plotted in Figure 9(a) for the logistic and Gompertz models. As before in Figure 8, the quantity 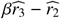 approaches zero as *β* becomes sufficiently small, while the quantity 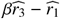 is approximately zero when *β* = 1, as expected. Plotting these quantities with noise-free data in Figure 9(a) illustrates how the MLE for 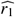 and 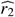 varies with *β* without the influence of external noise. Analogous data in Figure 9(b) shows the influence of corrupting the data with some noise and we see that the same underlying trends in the quantities 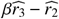 and 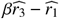 as functions of β are modestly impacted by incorporating noise in to the data.

**Figure 9:**
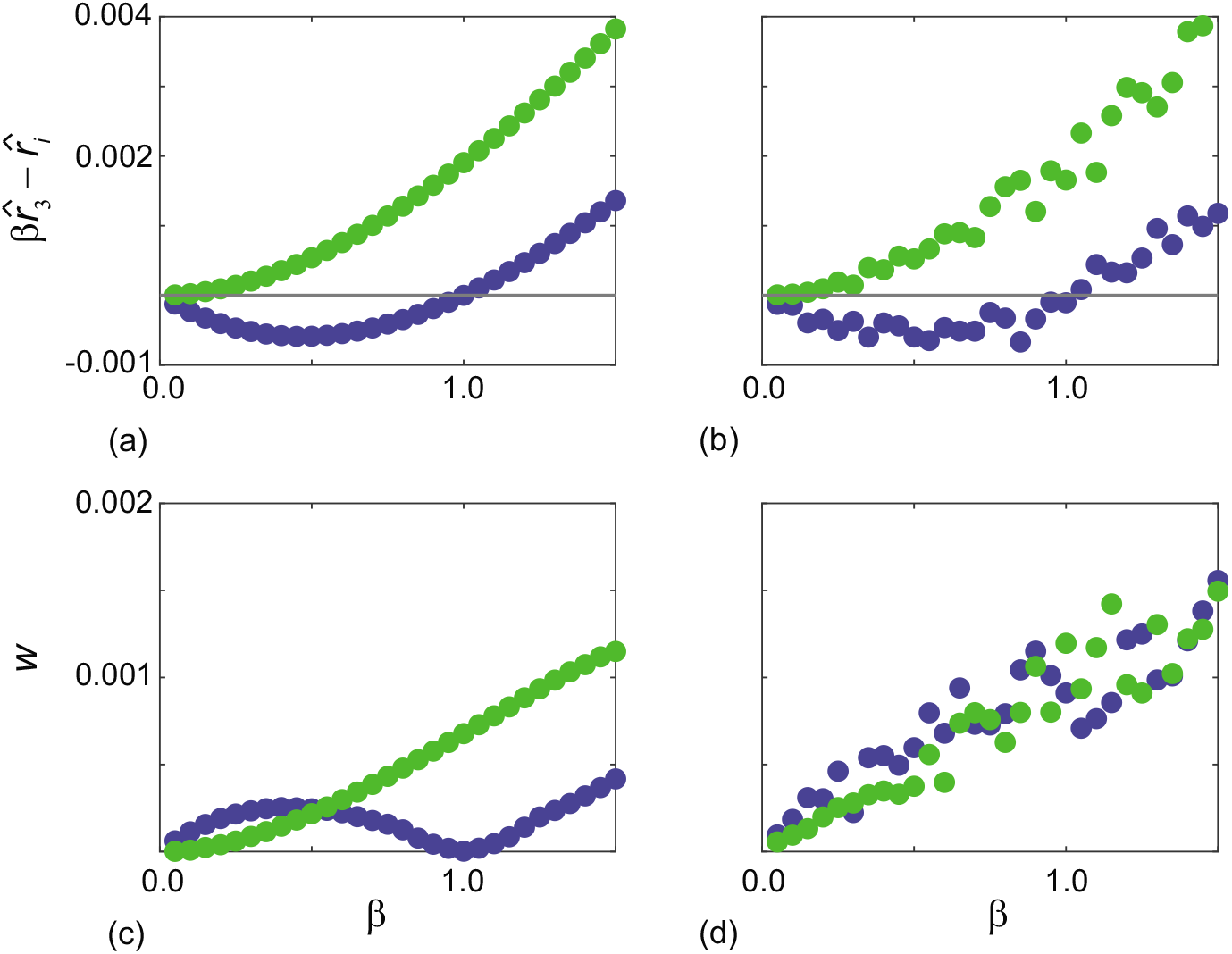
(a)–(b) show quantities 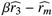 for *m* = 1, 2 as functions of *β* for noise free data (*σ* = 0) and noisy data (*σ* = 2), respectively. (c)–(d) show the width of the profiles, *w*, plotted as functions of *β* for noise free data (*σ* = 0) and noisy data (*σ* = 2), respectively. In all cases results for the logistic growth model are shown in blue while results for the Gompertz growth model are shown in green.

We now quantify the precision of the estimates of *r*_1_ and *r*_2_ in terms of the width of the 95% confidence interval, *w*. Data in Figure 9(c) shows how the width of the profiles varies with *β* for noise-free data, *σ* = 0, where we see the width of the profile for the logistic model approaches zero when *β* = 1, while the width of the profile for the Gompertz growth model approaches zero as *β* becomes sufficiently small, as expected. Additional results in Figure 9(d) show how the addition of noise to the data affects the width of the profiles, where we see that the clear trends in Figure 9(c) are completely obscured by the incorporation of noise.

## 3. Conclusion and Future Work

In this work we present a general framework that can aid model selection by comparing different models through the lens of practical parameter identifiability and checks of statistical assumptions underlying valid identifiability analysis. Population biology is one of many fields where questions of model selection are extremely important, with the potential to be very challenging because it is possible to interpret data with more than one candidate model [18]. In practice it is often unclear which model is most appropriate to work with and so developing methodologies that can assist in supporting these decisions is valuable.

Various approaches are adopted to deal with model selection in the literature, from the very simplest and surprisingly common approach of neglecting the question completely, to using more developed ideas such as information criteria [1, 4, 9, 25, 44, 56] and Bayesian approaches using Bayes factors [6, 26, 51]. Many of these established approaches, however, are difficult to interpret inferentially [13] and do not explicitly consider the fundamental question of practical identifiability where we assess whether the data contains sufficient information to form precise parameter estimates, and whether this varies between different candidate models.

For our particular data, we show that a simple sigmoid-growth predicted by several candidate mathematical models without any obvious challenges at first. Only when we ask the question whether the parameters are identifiable does it becomes clear that some of these models are very attractive while others are not. In this case we show that typical data relating to sigmoid re-growth of hard corals can be modelled using logistic, Gompertz and Richards’ models. However, both the Gompertz and Richards’ models encounter identifiability issues, whereas there is no such issue with the logistic model. While this finding is consistent with our intuitive expectation that increasingly complicated models ought to be used only when there is sufficient data available to estimate the unknown parameters, the difference in identifiability is somewhat surprising. For example, the logistic growth model with four unknown quantities *θ* = (*r*_1_, *K*, *C*(0), *σ*) does not lead to any identifiability issues, whereas the more commonly-invoked Gompertz growth model with the same number of unknowns, *θ* = (*r*_2_, *K*, *C*(0), *σ*), encounters practical identifiability issues since estimates of *C*(0) are not identifiable. This difference is surprising given that these two models involve the same number of unknowns, and even more surprising in this context since the Gompertz growth model is routinely used in the study of coral reef re-growth, but often this model is used without any explicit consideration of parameter identifiability [53, 57].

As we demonstrated, attempts to simplify parameter estimation by the common but ad-hoc practice fixing of initial conditions introduces separate issues with the statistical assumptions underlying valid identifiability analysis, as indicated by residual checks. This suggests that great care ought to be taken when we interpret experimental data with a mathematical model since the temptation that more complex models lead to better insight is not necessarily true. We suggest that the adoption of practical identifiability analysis along with model checks (e.g. inspection of residuals) is a very straightforward way to help guide model selection, and thereby aid in understanding underlying biological mechanisms. While our work here focuses on three simple models and one sigmoid data set, our approach is broad and applies to any continuum sigmoid model and any appropriate sigmoid data set. Further, our approach of selecting a model based upon the degree to which parameters are identifiable, provided it also passes statistical misspecification checks, has a strong implication for the question of experimental design, since we can use the tools presented in this work to explore what additional data is required to improve the identifiability of the parameters [55]. We leave the question of experimental design for future consideration.

## Acknowledgements

MJS is supported by the Australian Research Council (DP200100177). OJM received support from the University of Auckland, Faculty of Engineering James and Hazel D. Lord Emerging Faculty Fellowship. REB acknowledges the Royal Society for a Wolfson Research Merit Award, and the BBSRC (BB/R000816/1).

## Author Contributions

All authors conceived ideas the designed methodology, MJS developed software and led the writing of the manuscript. All authors analysed the data and contributed critically to the drafts and gave final approval for publication.

## Data Availability

Data and software are available on GitHub, and the complete LTMP data set is available at [eAtlas](https://eatlas.org.au/gbr/ltmp-data).

## Notes

https://github.com/ProfMJSimpson/SigmoidGrowth

### Competing Interest Statement

The authors have declared no competing interest.

